# EEG-guided robotic control of transcranial magnetic stimulation

**DOI:** 10.64898/2026.05.15.725366

**Authors:** Renan H. Matsuda, Matilda Makkonen, Ivan Zubarev, Olli-Pekka Kahilakoski, Leevi A. Kinnunen, Mila Nurminen, Simo-Pekka Simonaho, Risto J. Ilmoniemi, Victor H. Souza, Pantelis Lioumis

## Abstract

Brain-state-guided and closed-loop transcranial magnetic stimulation (TMS) protocols have emerged as methods for decreasing the outcome variability and increasing the therapeutic effectiveness of stimulation protocols. However, existing brain-state-dependent TMS systems control only stimulation timing, while the location is fixed and manually adjusted between blocks or sessions. This limits targeting distributed cortical networks in a flexible brain-state-dependent paradigm.

We developed a method that jointly manages TMS pulse timing and location automatically controlled by electroencephalography (EEG)-derived brain state. A machine-learning algorithm infers the brain state in real time to guide robotic coil placement. We present a proof-of-concept study in which EEG controlled both the target site and the timing of TMS. A pre-trained convolutional neural network discriminated between resting state and movements performed with the right or left hand; the classifier output determined the hemisphere in which primary-motor-cortex hand area was stimulated. Preprocessing and decoding of 2-s EEG segments required 150 ms, and the robot took 7.5 s to move from the vertex home position to the predefined motor targets.

The EEG-guided robotic TMS method expands the toolkit for brain-state-dependent neurostimulation by enabling closed-loop control of stimulus location based on EEG-derived brain states, including volitional activity. Thus, the method can benefit both neuroscience research and clinical neuromodulation applications, such as by optimizing stimulation parameters to the brain state that produces optimal rehabilitation outcomes in spinal cord injury or motor disorders.

## 1. Introduction

Transcranial magnetic stimulation (TMS) is a non-invasive neuromodulation method that employs brief, intense magnetic pulses to induce electric fields in the cortex, depolarizing neuronal membranes (Barker et al., 1985). TMS assisted by neuronavigation (Hannula and Ilmoniemi, 2017) has shown great potential for motor cortical mapping (Picht et al., 2009; Vitikainen et al., 2009) due to the spatial specificity and the possibility to target individual muscle groups. TMS is also used for speech cortical mapping (Lioumis et al., 2012) as a presurgical evaluation prior to neurosurgery (Krieg et al., 2014), for motor rehabilitation (Shulga et al., 2021), and mood disorder treatments. Conventional TMS requires manual placement of the coil above the stimulation site. However, combined with a robotic arm, the stimulation location can be autonomously moved between cortical areas without any manual involvement (Matsuda et al., 2024).

Simultaneous TMS and electroencephalography (TMS–EEG) allows perturbing and measuring brain function; focal TMS enables the stimulation of specific neural populations, whereas the high temporal resolution of EEG provides an instantaneous view of the responses evoked by TMS (Ilmoniemi et al., 1997). Previous research has demonstrated that brain stimulation during different brain states has different modulatory effects (Silvanto et al., 2008; Zrenner et al., 2023, 2018), indicating that EEG-defined brain-state-guided TMS could be therapeutically more effective than current protocols (Zrenner and Ziemann, 2024). Brain-state-guided stimulation requires real-time EEG analysis with the help of algorithms to categorize EEG signal patterns.

Existing protocols for brain-state-dependent stimulation focus on stimulation timing control based on EEG oscillation features, such as mu-rhythm phase (Zrenner et al., 2023, 2018). This technology is limited to one stimulation location with a fixed coil placement. However, cognitive and behavioral brain functions are known to operate in distributed brain networks instead of singular cortical locations (Gordon et al., 2023; Lynn and Bassett, 2019; Siddiqi et al., 2022; Yang et al., 2021). In order to probe and interact with these functions optimally, we need to engage with the dynamics of distributed networks. Stimulating multiple nodes in a network based on brain state might uncover new findings on causal communication between nodes, helping to understand the mechanisms behind brain network function (Siddiqi et al., 2021). To our knowledge, brain-state-dependent TMS systems that control both stimulation timing and location have not been presented. Here, we take advantage of a robotically placed TMS system (Matsuda et al., 2025), which broadens the possibilities for brain-state-dependent stimulation.

Brain-state-dependent stimulation can be expanded into closed-loop paradigms, in which parameters are changed adaptively based on feedback from TMS-evoked EEG potentials (TEPs; Bergmann, 2018; Zrenner and Ziemann, 2024). Closed-loop EEG–TMS protocols offer promise in personalized treatment (Humaidan et al., 2024), as they enable real-time optimization of temporal and spatial targeting. Since the required protocols and algorithms are complex, few reports of fully closed-loop EEG–TMS studies have been published (Humaidan et al., 2024; Tervo et al., 2022).

Brain–computer interfaces (BCIs) are another important application area of real-time brain-state detection. BCIs have attained a great deal of interest due to their potential in the treatment or rehabilitation of patients suffering from severe motor disabilities (Lebedev and Nicolelis, 2017; Teo and Chew, 2014). Specifically, BCI systems combined with motor imagery (MI) can translate the participant’s brain activity associated with imagined limb movements into commands to control an external device (Padfield et al., 2019). In MI-BCI, when the user imagines moving a limb, oscillatory events are observed in EEG signals originating from brain areas associated with the preparation, control, and execution of voluntary motion (Pelgrims et al., 2011; Schnitzler et al., 1997). Moreover, MI shows potential to augment motor rehabilitation (Liu et al., 2023; Pichiorri et al., 2015; Ramos-Murguialday et al., 2019; Ramos-Murguialday et al., 2013; Teo and Chew, 2014). TMS during MI has been shown to facilitate motor learning and induce cortical plastic changes (Chen et al., 2024; Foysal and Baker, 2020; Muraoka et al., 2024; Kraus et al., 2016). In current rehabilitation paradigms, the extent, quality, and timing of MI in individual patients are unclear, which may increase the variability of results. Combining the TMS rehabilitation protocol with MI-BCI may result in more precisely timed and efficient stimulation (Kraus et al., 2016; Royter and Gharabaghi, 2016; Jackson et al., 2001), while facilitating automation and potentially supporting home-based treatments.

Our aim was to develop a brain-state-dependent stimulation method in which EEG-measured brain activity guides the robotic TMS placement and controls pulse delivery. We demonstrate the method with a proof-of-concept experiment in which a motor-execution BCI automatically selects the cortical target and initiates robotic coil positioning followed by TMS pulse delivery. The presented method enables a wide range of closed-loop and brain-state-dependent EEG–TMS experiments and interventions, with fully automatic stimulation, flexible real-time EEG preprocessing and analysis, and synchronized visual stimulus presentation to the participant. The method can be used to develop treatment and rehabilitation protocols for personalized TMS interventions, in which stimulation timing and location are automatically controlled in real time based on spontaneous or volitional cortical responses to TMS and visual stimuli.

## 2. Technical setup

We developed a brain-state-dependent stimulation method in which EEG guides the robotic placement and timing of TMS. Our setup (Fig. 1) consists of an EEG and electromyography (EMG) device, a computer running real-time EEG processing and TMS control software, another computer running neuronavigation software connected to infrared cameras, a TMS stimulator and coil, and a robotic arm. The method builds on previously described software and robotic targeting components (Kahilakoski et al., 2025; Matsuda et al., 2025, 2024; Souza et al., 2018). Here, we developed a communication interface that transfers EEG-derived target selections to the neuronavigation software, enabling automatic switching between predefined stimulation sites, as well as a novel robotic control architecture that regulates coil–scalp contact during positioning. We integrated these components into an automated workflow that coordinates EEG-based target selection, visual cue presentation, and navigated robotic coil positioning with TMS pulse delivery enabled once the positioning criteria are satisfied.

**Figure 1:**
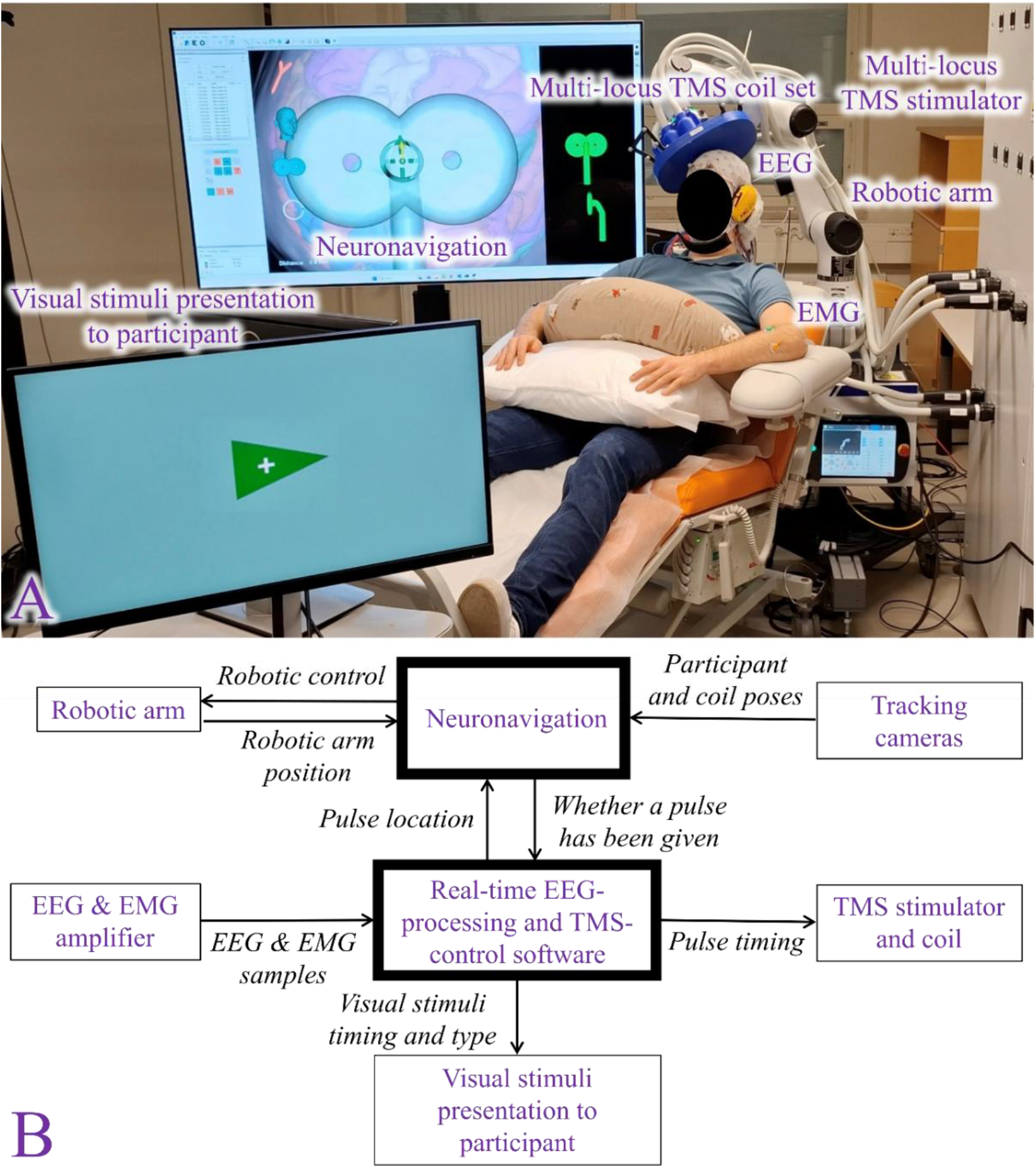
Components and information flow in the EEG-controlled TMS system. Panel A: Experimental setup with key hardware annotated: multi-locus TMS coil set (Nieminen et al., 2022) mounted on a robotic arm, multi-locus TMS stimulator, EEG cap, EMG electrodes, neuronavigation system, and the monitor for presenting visual stimuli to the participant. The neuronavigation screen shows the individual head model and current coil position (see Supplementary Video). Panel B: Block diagram of data and control flow. EEG and EMG signals are acquired by the amplifier and streamed to the real-time EEG-processing and TMS-control software, which also sends information on visual-stimulus timing and type. Based on the EEG and EMG inputs, the software determines pulse timing and location and sends commands to the TMS stimulator and neuronavigation. Neuronavigation tracks the positions of the participant and TMS coil set with tracking cameras and controls the robotic arm position, enabling closed-loop robotic control of coil pose.

We set a criterion that the method must meet for successful performance. The robotically guided coil must reach its desired spatial target even when moving between hemispheres. We quantify this with physiological output by measuring the motor-evoked potentials (MEPs) from primary motor cortex stimulation; obtaining an MEP from each pulse confirms the reliability of automatic coil positioning.

### 2.1. General workflow

We implemented a method that reads ongoing brain activity, decodes it, moves the TMS coil robotically to the desired location based on the decoded brain activity, and delivers TMS pulses at the desired time. The technical workflow is as follows (Fig. 1B): The real-time EEG-processing software performs EEG/EMG analysis (*e.g.*, classification) and selects the stimulation target according to the analysis results. The target is then sent to neuronavigation. The neuronavigation software represents the selected target as a six-degree-of-freedom (DOF) coil pose (position and orientation), defined relative to the local scalp surface, using the surface normal at that point and a tangential direction along the scalp. Using real-time estimates from the tracking cameras, the neuronavigation software computes the instantaneous displacement between the current coil pose and the target pose and continuously sends this displacement to the robot controller, which updates the coil position until both pose and contact-pressure tolerances are satisfied. When these criteria are met, the neuronavigation software asserts the stimulation-enable gate. The real-time EEG-processing software then sends a time-stamped trigger to the TMS controller, which delivers the TMS pulse and returns a marker to the EEG amplifier to register the exact pulse timing.

### 2.2. Real-time EEG-processing software

In this work, we developed communication between EEG processing and neuronavigation to enable EEG-based stimulation location changes. For this, we utilized the NeuroSimo software (Kahilakoski et al., 2025), which is used for real-time EEG and EMG processing as well as TMS control. NeuroSimo is responsible for receiving, processing, and analyzing EEG and EMG data streamed from the amplifier, and integrating them with stimulation timing decisions and visual stimuli presented to the participant. In this study, we implemented communication between NeuroSimo and the neuronavigation software to translate EEG-classification results into stimulation-target requests.

The software comprises three main modules, forming the data processing pipeline: Preprocessor, Decider, and Presenter. Processing within each module is implemented as a Python script, while the backend of the software handles the lower-level operations, such as preparing data buffers. The Preprocessor module performs sample-by-sample EEG preprocessing, such as spatial filtering and moving-window baseline removal, and sends each preprocessed sample forward to the Decider. The Preprocessor is important for EEG–TMS, as it allows removing noise from the EEG in a sample-by-sample manner before decoding brain states, thus increasing decoding performance without introducing processing latencies.

The Decider module classifies EEG patterns, makes decisions, and communicates them to the TMS controller and the neuronavigation software. The module backend accumulates EEG signals into buffers, allowing the Python script to preprocess them and classify the resulting data with a variety of algorithms. Algorithm training during an experiment is also possible. Data buffers can be preprocessed with MNE-Python (Gramfort et al., 2013), for example, by applying bandpass filters to isolate relevant EEG frequency bands. The Decider can also analyze EMG signals, such as MEPs elicited by TMS.

NeuroSimo’s Decider module uses predefined state-machine logic to translate EEG-classification results into stimulation decisions. For this study, we implemented a communication interface that sends the selected stimulation target from NeuroSimo to the neuronavigation software. Neuronavigation then guides robotic coil positioning and enables stimulation once the positioning criteria are satisfied. NeuroSimo coordinates pulse timing and delivery through the TMS controller. This sequence links EEG-based selection of predefined targets to robotic coil positioning and subsequent stimulation.

The Presenter module utilizes PsychoPy (Peirce, 2007) to present visual and auditory stimuli to the participant. It can dynamically adapt inter-trial intervals or stimulus types based on EEG-classification results, supporting experiments where sensory stimuli are adjusted according to brain-state analysis. Adaptive sensory stimulus delivery enables a variety of BCI protocols, increasing the versatility of the EEG-guided TMS method.

In this study, we developed experiment-specific NeuroSimo Decider and Presenter scripts, which are available at https://github.com/connect2brain/bci-tms. The NeuroSimo software itself is available at https://github.com/NeuroSimo.

### 2.3. Neuronavigation software and tracking cameras

Neuronavigation is performed with the InVesalius software, whose development and characterization have been reported previously (Souza et al., 2018). InVesalius is used to reconstruct participant-specific 3D anatomical models from structural MRI and to define the cortical target sites for stimulation. An OptiTrack system with eight Flex13 infrared cameras tracks the participant’s head and the TMS coil. This ensures spatial precision and reproducibility of stimulation with respect to individual cortical anatomy, which is important in brain-state-dependent and closed-loop TMS experiments with multiple adaptively switched stimulation targets.

For this study, we extended the InVesalius communication interface to receive stimulation-target requests from NeuroSimo. Each request specified a target label that InVesalius matched to a predefined coil target containing the desired position and orientation. The target-update routine temporarily stopped robotic target tracking, selected the requested target, and reactivated tracking for the new target. This extension enabled EEG-driven switches between predefined stimulation sites without manual target selection in the neuronavigation interface.

During navigation, InVesalius continuously computed the displacement between the current coil pose and the selected target pose and supplied this information to the robotic controller. Stimulation was enabled only when the three-dimensional Euclidean distance between the tracked coil position and the target position was less than or equal to 2 mm and the absolute value of each of the three Euler-angle components of the relative rotation was less than 2°. All four conditions had to be satisfied simultaneously. The neuronavigation software transmitted head and coil poses, tracking status, and the selected target to the robot controller to guide coil positioning and support safety checks. The controller stopped motion if tracking was lost or head velocity exceeded the configured limit, and rejected targets outside the robot workspace. These safeguards are particularly important when stimulation targets are automatically, adaptively, and rapidly changed during an experiment.

### 2.4. Robotic control

In this study, we implemented a new control architecture that builds on our earlier work in robotic TMS positioning and force control (Matsuda et al., 2025, 2024). Robotic positioning is performed with an Elfin E5 six-DOF collaborative arm, commanded by neuronavigation. Control is formulated as a multi-axis proportional-integral-derivative (PID)/impedance scheme driven by the displacement in coil-set position and orientation provided by neuronavigation.

At each control cycle (Fig. 2), translation components (*dx*, *dy*, *dz*) and rotation components (*drx*, *dry*, *drz*) update axis-specific controllers. Lateral (*x*, *y*) and rotational (*rx*, *ry*, *rz*) axes use displacement-based PID control to reduce pose error relative to the target. These features ensure quick coil set movement between distant cortical targets, which facilitates and accelerates all TMS experiments with stimulation sites in multiple brain areas.

**Figure 2:**
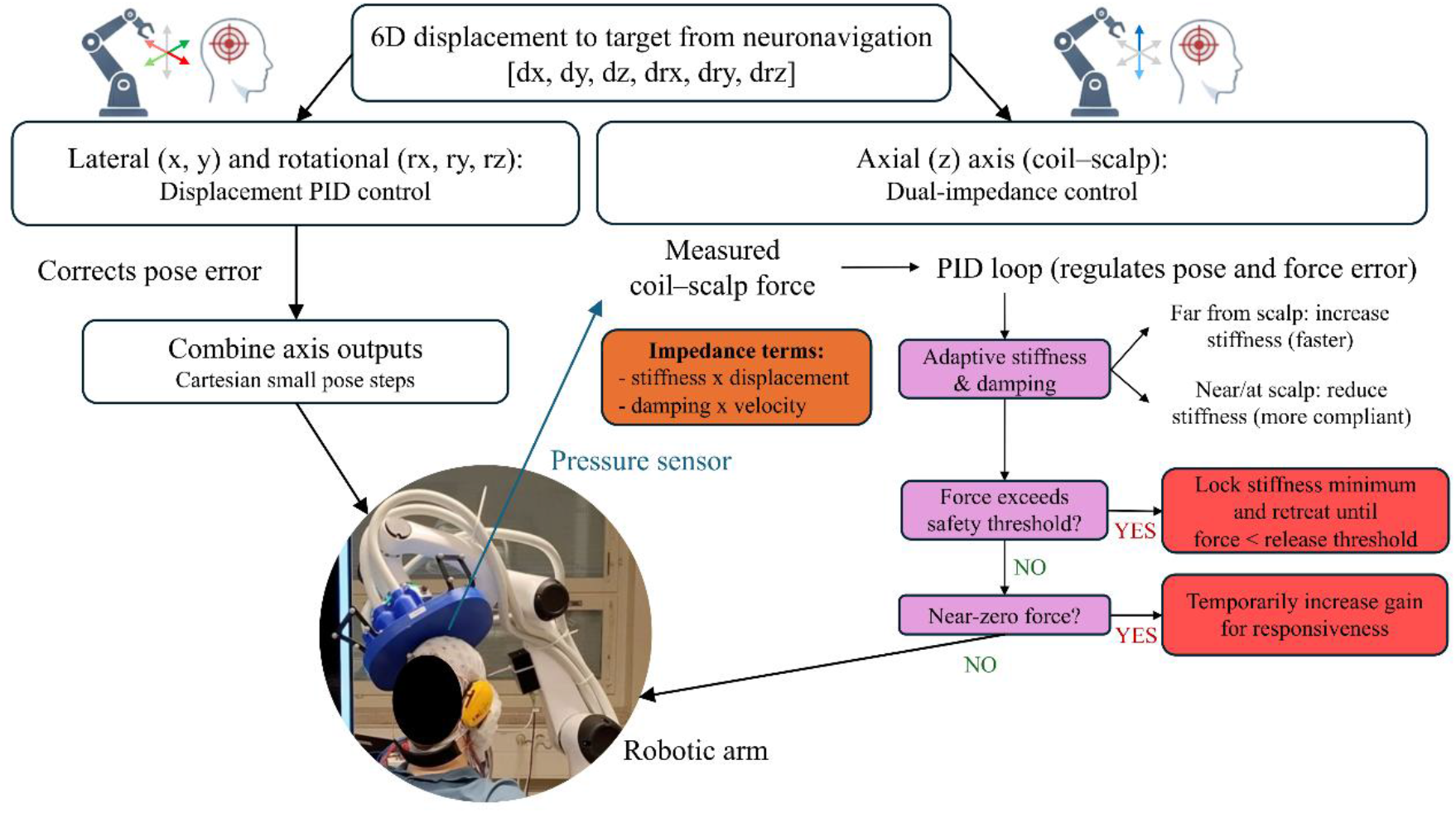
Control architecture for robotic TMS coil positioning. A 6D displacement-to-target vector from neuronavigation (translation: *dx, dy, dz*; rotation: *drx, dry, drz*) drives displacement proportional-integral-derivative (PID) control of the lateral (*x, y*) and rotational axes to correct pose displacement, while the axial (*z*) axis uses dual-impedance control based on coil–scalp distance and force from a pressure sensor. Axis outputs are combined into Cartesian small pose increments to command the robotic arm. An adaptive stiffness–damping loop regulates force displacement in *z*, increasing stiffness when far from the scalp, reducing stiffness near contact, and triggering safety behaviors (stiffness lock, pause/retreat, or temporary gain increase) when force thresholds are exceeded or near zero. This architecture provides regulated coil–scalp contact, enabling participant comfort and safety, reproducibility, and precision of the stimulation location.

The axial (*z*) axis uses a dual-impedance controller with a pressure sensor integrated at the coil-set–scalp interface (Kim and Yang, 2024). In this mode, a PID loop regulates force displacement relative to a desired contact force setpoint, while impedance terms (stiffness times displacement and damping times velocity) provide compliant behavior. The controller adapts stiffness and damping in real time based on the *z* displacement and measured force: stiffness is increased when the coil set is further from the target location to accelerate approach and reduced near or at contact to enhance compliance. If the measured force exceeds a safety threshold, stiffness is locked to a minimum and the controller either pauses or retreats until the force decays below a release threshold. If the coil set is not in contact with the scalp, proportional gains are temporarily elevated for responsiveness during initial contact acquisition and reduced under load to maintain stability. This design maintains a gentle and regulated contact between the TMS coil set and the scalp, ensuring participant comfort and safety as well as reliability and precision of the stimulation location. These qualities are required in closed-loop TMS experiments with changing stimulation locations. The open-source implementation of the robotic arm control is available at https://github.com/biomaglab/tms-robot-control.

### 2.5. TMS hardware

Our method can be used with any TMS coil attached to a robotic arm. Each state decoded by BCI can be mapped to a predefined cortical target within the robotic TMS workspace. In a proof-of-concept experiment, we used the novel mTMS system for convenience, since it is software-controllable by design. The mTMS system can electronically steer the induced electric field on a millisecond level without repositioning the coil set (Nieminen et al., 2022). However, we did not use this electric-field steering in the presented method. Electronic steering can be used for brain-state-dependent control of stimulation location because it allows millisecond-level, programmable changes in the cortical target driven directly by the BCI algorithm.

## 3. Proof-of-concept experiment

We demonstrated the functionality of the EEG-controlled TMS method with a proof-of-concept experiment involving motor-execution (ME)-BCI. The purpose of the experiment was to demonstrate the integration of the novel components and show that EEG decoding could be used to automatically control the subsequent stimulation sequence and target selection. The experiment allowed measuring whether the method fulfilled the criterion regarding spatial targeting accuracy.

A 22-year-old right-handed male with no neurological or motor disorders participated in the experiment. The study was accepted by the Coordinating Ethics Committee of the Hospital District of Helsinki and Uusimaa (HUS/1198/2016), and it followed the Declaration of Helsinki. The participant provided informed consent prior to the study.

The participant was instructed to move either the left or right hand, or to perform no movement. The movement type was classified from the EEG with a machine-learning algorithm. If the EEG was classified as a left-hand movement, the mTMS coil set was robotically moved, and a pulse was delivered to the right primary motor cortex (M1), and vice versa. If no movement was detected, no stimulation was given. The details of the experiment are provided below.

### 3.1. Training data

The data for training the ME-classification algorithm were collected immediately prior to the TMS experiment. EEG data were obtained at a sampling rate of 1 kHz with a Bittium NeurOne amplifier and a TMS-compatible 62-channel cap. Electrode impedances were kept below 5 kΩ.

The participant was visually cued via NeuroSimo Presenter to execute one of three trial types: move the left or right hand or move nothing. The hand movement was defined as a wrist extension, and while not executing movements, the participant kept his hands resting on a pillow in his lap (see Fig. 1A). The participant was instructed to avoid other movements.

The data were collected in 20 blocks of 30 trials each. Each block included 10 trials of each trial type, in randomized order. Each trial consisted of a 1-s visual instruction, followed by 1.5 s of movement execution. The inter-trial interval was randomized from a uniform distribution of 1–3 s. At the beginning of each trial, NeuroSimo sent a trigger marker to the EEG through a LabJack T4 USB peripheral and saved the trial timestamps and types in a log file. After each block, there was a 20-s break. Additionally, three 2-minute breaks were included after the 5^th^, 10^th^, and 15^th^ blocks.

The participant viewed the visual cues on a screen. A fixation cross was always shown except during inter-trial intervals. The visual instruction for movement was a green arrow overlaid on the fixation cross, pointing in the direction of movement (Fig. 1A). The participant was instructed to begin the movement after the arrow disappeared. The non-movement trial instruction was a yellow hourglass shape superimposed with the fixation cross.

### 3.2. Algorithm training

The ME-detection algorithm was trained during a short break between the training data collection and the TMS session. EEG data were first epoched from 0–2 s after each movement-start trigger. Epochs were then bandpass filtered between 0.1 and 70 Hz and transformed to the common average reference. Each epoch was also baseline-corrected by removing the mean of the epoch for each channel and scaled by dividing each value in the epoch by the standard deviation across all channels and time points. Each epoch was labeled as “left”, “right”, or “no movement” based on the NeuroSimo log file. Preprocessing was performed with MNE-Python (Gramfort et al., 2013).

We chose the VAR-CNN (Zubarev et al., 2019) algorithm for ME classification, as it was developed for magnetoencephalography (MEG)/EEG data classification and validated on MI data. The preprocessed epochs were divided into 5 folds, and training was performed in a cross-validation setup using the MNEflow software package (Zubarev et al., 2022) with parameters given in Table 1.

**Table 1:**
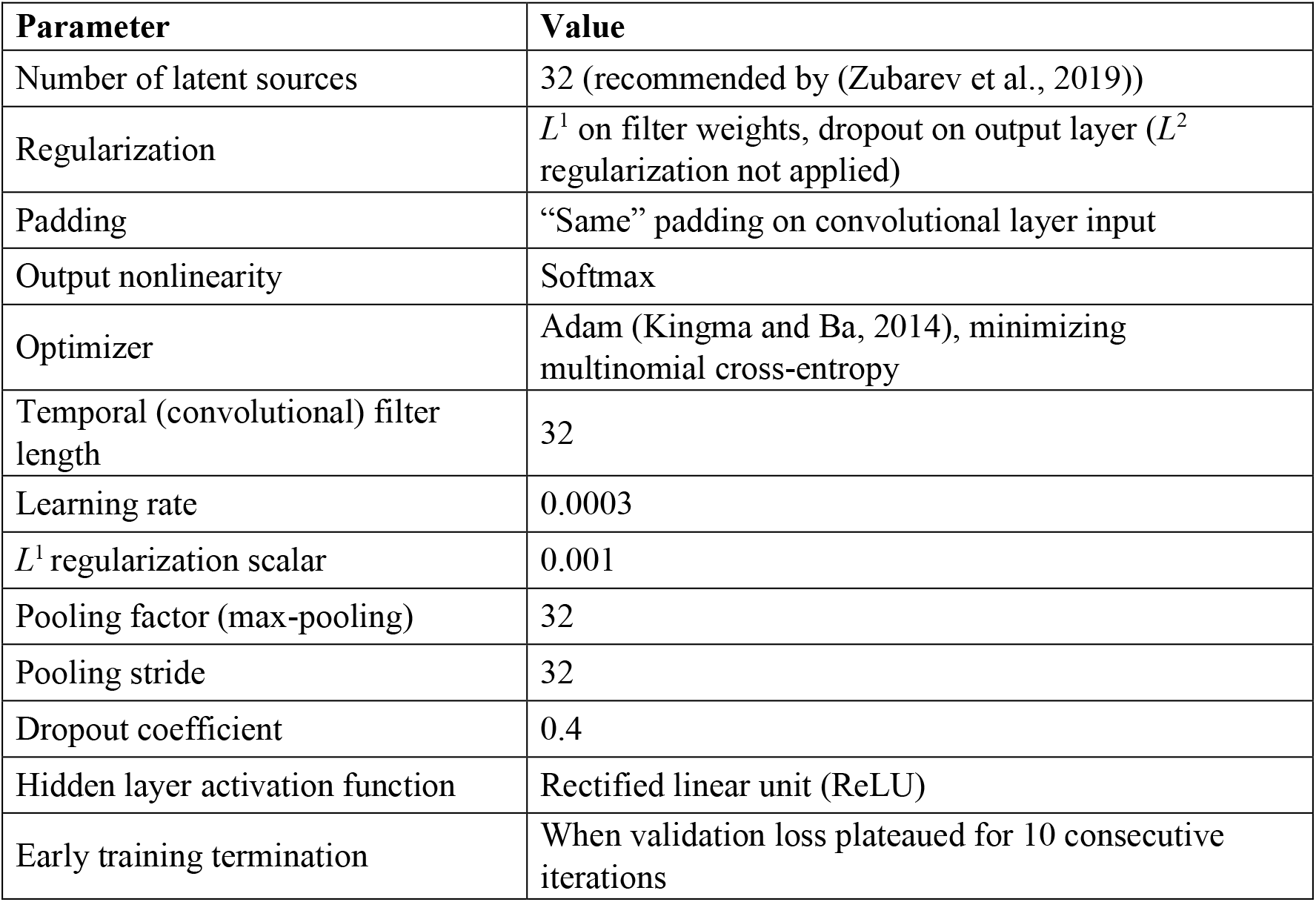
Architecture parameters and hyperparameter values for ME-classification machine-learning algorithm VAR-CNN training.

| Parameter | Value |
| --- | --- |
| Number of latent sources | 32 (recommended by (Zubarev et al., 2019)) |
| Regularization | $L^1$ on filter weights, dropout on output layer ($L^2$ regularization not applied) |
| Padding | “Same” padding on convolutional layer input |
| Output nonlinearity | Softmax |
| Optimizer | Adam (Kingma and Ba, 2014), minimizing multinomial cross-entropy |
| Temporal (convolutional) filter length | 32 |
| Learning rate | 0.0003 |
| $L^1$ regularization scalar | 0.001 |
| Pooling factor (max-pooling) | 32 |
| Pooling stride | 32 |
| Dropout coefficient | 0.4 |
| Hidden layer activation function | Rectified linear unit (ReLU) |
| Early training termination | When validation loss plateaued for 10 consecutive iterations |

### 3.3. Real-time EEG–TMS experiment

In the experiment (Fig. 1A and Supplementary Video), the participant was instructed to perform three consecutive cued repetitions of the same trial type (left or right wrist extension or no movement). The EEG signal from each trial was classified in real time using the trained algorithm. The final classification was determined by the majority vote from the three repetitions; if all three were classified into different types, the final classification was defined as “no movement”. Depending on the final classification, the mTMS coil set was automatically moved by the robotic arm to stimulate the M1 of the corresponding hemisphere.

Before starting the real-time experiment, the left and right M1 hotspots for the *extensor carpi radialis* muscles (responsible for wrist extension) were determined and defined as the LEFT-M1 and RIGHT-M1 targets. We determined the hotspots with the novel automatic BOOST hotspot-hunting algorithm (Granö et al., 2025; Tervo et al., 2020). Electrodes were placed on these muscles in the belly-tendon montage to record EMG using the Bittium NeurOne amplifier. The stimulation intensity was set to produce a cortical electric field of 100 V/m, which elicited MEPs at both left and right M1 hotspots. After stimulation, the robotic arm returned the coil set to the predefined HOME position over the participant’s vertex, *ca.* 10 cm above the scalp (see Supplementary Video).

The visual cues presented to the participant in the real-time experiment were the same as during training data collection. Sample-by-sample preprocessing with the Preprocessor module was not utilized in the proof-of-concept experiment. Immediately following trial end, a buffer consisting of the trial data was preprocessed with the NeuroSimo Decider module identically to the algorithm training data (see Section 3.2) and classified with the pre-trained VAR-CNN algorithm using Tensorflow (Abadi, 2016). After three consecutive same-movement trials, the majority vote of the classifier outputs decided the stimulation target: “left” classification mapped to RIGHT-M1, “right” to LEFT-M1, and “no movement” to the HOME position, in which no stimulation was delivered. The majority-vote classification result was visually presented to the participant via NeuroSimo Presenter.

NeuroSimo communicated the stimulation decision request to the neuronavigation software. Neuronavigation checked spatial and safety criteria, setting the stimulation-enable gate if conditions were met. The robotic arm moved the mTMS coil set to the target (Fig. 2); once positioned, the TMS controller delivered a pulse. The sets of three continuous motor executions were repeated 30 times: 10 times for each trial type.

### 3.4. Analysis of experiment results

To evaluate method performance according to the pre-set criterion, we extracted MEPs for LEFT-M1 and RIGHT-M1 stimulation to verify coil-set positioning reliability across robotic arm movements. The continuous EMG data were bandpass filtered between 20 and 450 Hz to remove slow drifts and artifacts unrelated to the MEP, and notch-filtered to remove the 50-Hz line noise. MEP peak-to-peak amplitudes were calculated from the preprocessed EMG as the difference between the largest and smallest signal value within the time window 10–60 ms after TMS pulse. Additionally, we calculated two latency metrics: the time from trial end to classification result (including trial preprocessing and decoding), and the time from initiation of robotic arm movement from the HOME position to pulse delivery. We conducted a one-way ANOVA to compare classifier latencies across trial types.

Lastly, we analyzed accuracy and precision of the majority-vote EEG-classification (see Section 3.3). The accuracy was calculated as the proportion of correct classifications, *i.e.*, where classification output matched the condition cued to the participant, out of all classifications. The precision was calculated separately for each of the three conditions as the number of the correctly classified majority-vote results (*i.e.*, those matching the condition cued to the participant) divided by the total number of majority-vote results assigned to that condition.

## 4. Results

The EEG-guided TMS method fulfilled the pre-set criterion for spatial targeting accuracy. MEPs were elicited in response to all TMS pulses to both stimulation targets (Fig. 3), indicating reliable positioning over the intended cortical sites. The results below are presented as mean ± standard deviation. LEFT-M1 stimulation elicited MEPs in the right *extensor carpi radialis* muscle with a peak-to-peak amplitude of 1300 ± 500 µV (*N* = 6), while stimulation to RIGHT-M1 elicited MEPs in the left *extensor carpi radialis* muscle with an amplitude of 1000 ± 560 µV (*N* = 15).

**Figure 3:**
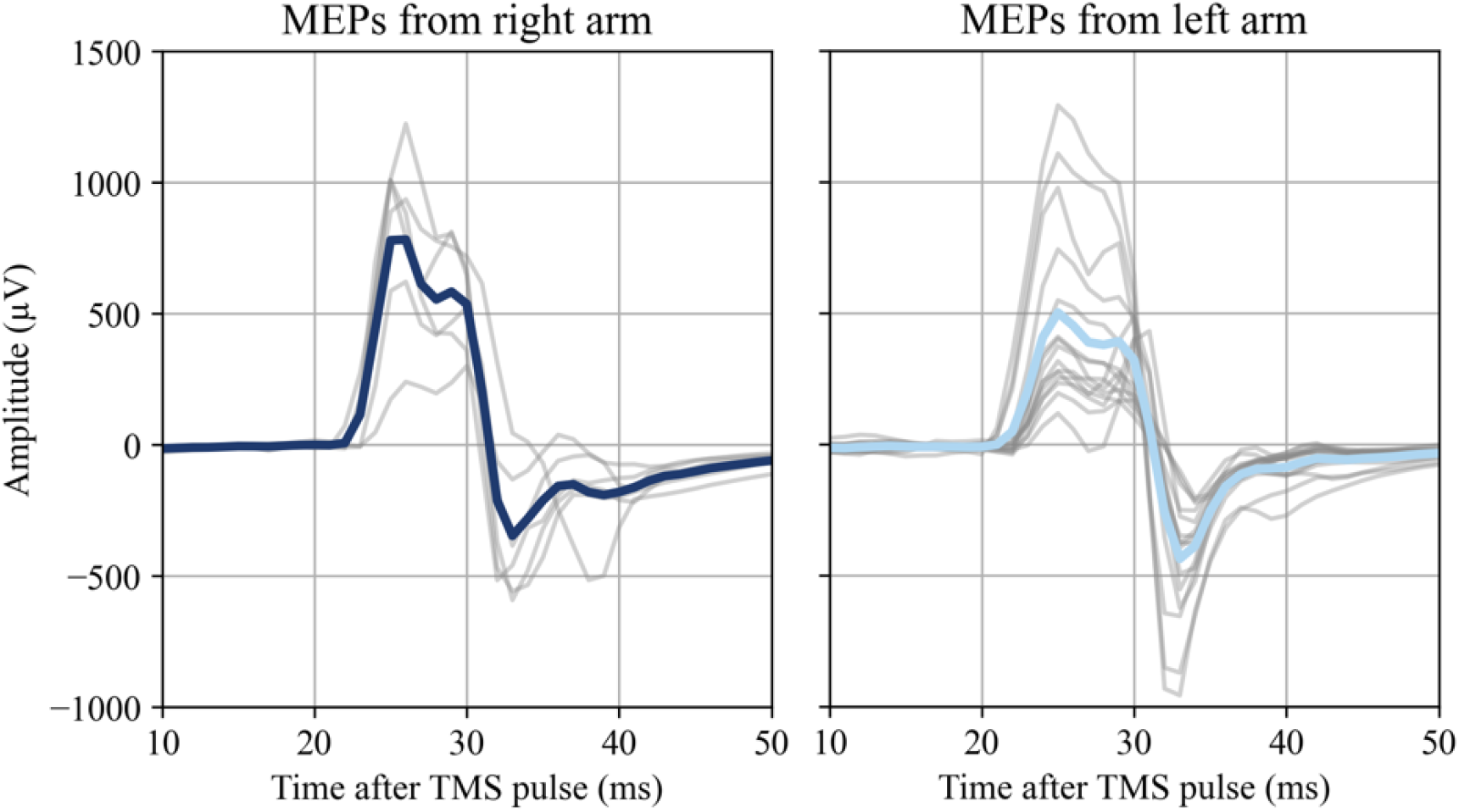
Motor-evoked potentials (MEPs) measured from the *extensor carpi radialis* muscles from the two stimulation sites: LEFT-M1 for right-arm MEPs (*N*=6) and RIGHT-M1 for left-arm MEPs (*N*=15). The colored lines show the average MEPs and light grey lines show single-trial MEPs.

The classification latency across all trials was 150 ± 20 ms. The latencies were similar for trials decoded as left (140 ± 10 ms), right (150 ± 10 ms), and non-movement (150 ± 30 ms) (Fig. 4A). ANOVA revealed no significant differences in classification latencies across trial types (*F* = 0.59, *p* = 0.56), suggesting that the EEG-processing duration is stable.

**Figure 4:**
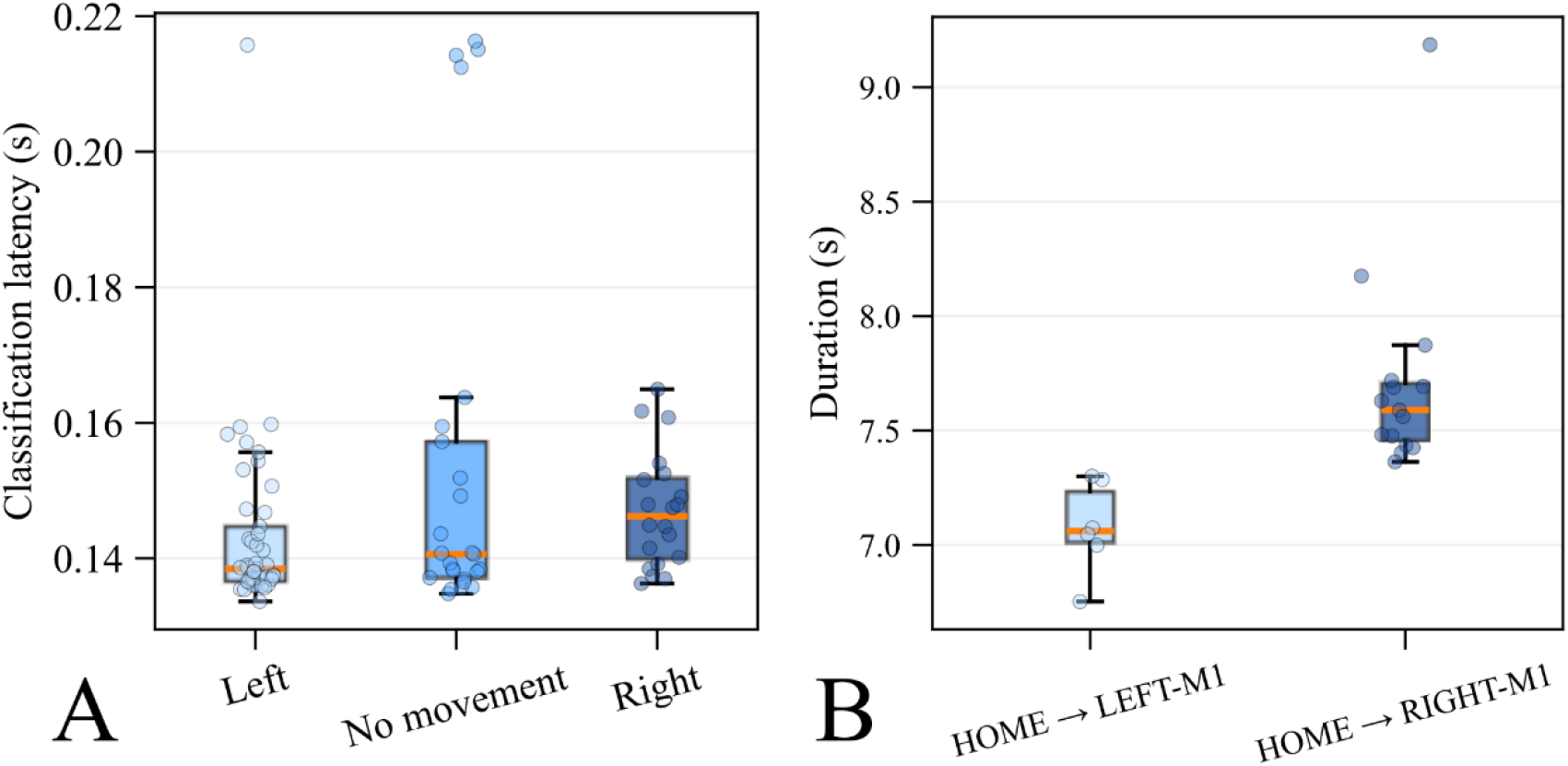
Panel A: Classification latencies (including preprocessing and decoding) of EEG trials by classification result. *N*=45 for trials classified as left, *N*=25 for no movement, and *N*=20 for right. Panel B: Robot movement latencies from initiating robotic arm movement in the HOME position above the vertex to delivering a TMS pulse according to target site. *N*=6 for LEFT-M1 and *N*=15 for RIGHT-M1.

The latency from initiating robotic arm movement to delivering the pulse was 7.5 ± 0.5 s. The latency for LEFT-M1 was 7.1 ± 0.2 s, slightly shorter than the latency for RIGHT-M1 at 7.7 ± 0.4 s (Fig. 4B). The minimum latency was 6.8 s, and the maximum latency was 9.2 s. The longer latency values are from trials in which robotic repositioning required additional safety checks or adjustments, often in response to small participant movements that required minor trajectory corrections before pulse delivery.

The ME classification algorithm achieved an overall accuracy of 70% at the block level (best-of-three decisions; 21 correct out of 30 blocks). Classification performance was highest for left blocks, all of which were correctly identified (10/10). In contrast, accuracy was lower for no-movement (6/10) and right (5/10) blocks. The algorithm classified 15 blocks as left, of which 10 were truly left (precision 67%). The algorithm classified 9 blocks as no-movement, 6 of which were truly no-movement (precision 67%), and classified 6 blocks as right, 5 of which were truly right (precision 83%).

## 5. Discussion

We developed a method for brain-state-dependent robotic TMS with EEG-guided stimulation location control. Through a proof-of-concept experiment, we demonstrated that the method preprocesses and decodes EEG with stable latency and consistently delivers stimulation with high spatial precision via robotically placed TMS.

### 5.1. Comparison with existing systems

Previous brain-state-dependent TMS studies have used EEG oscillatory features to determine stimulation timing (Humaidan et al., 2024; Wischnewski et al., 2024; Zrenner et al., 2018). Our method adds the important dimension of stimulation location, enabling less than 10-s location changes using robotically positioned TMS. The method extends programmable control to selecting spatially separated targets, demonstrating automatic execution of EEG-derived target decisions with stimulation conditional on positioning criteria. This aspect enables brain-state-dependent experiments targeting different network nodes.

The presented method allows adjusting stimulation timing in a brain-state-dependent manner. Although in the proof-of-concept experiment we applied stimulation conditional to the robotically controlled coil reaching the target, NeuroSimo allows our method to deliver stimulation with a consistent latency of a few milliseconds if the coil is already on target (Kahilakoski et al., 2025). In addition, the EEG processing in our method can operate with EEG-oscillation analysis (Kahilakoski et al., 2025) but is not limited to it. NeuroSimo flexibly operates with any brain-state-analysis algorithm that can be implemented in Python, including complex machine-learning algorithms. Thus, users can choose or develop an algorithm best suited to their needs. Additionally, our method includes sensory stimulus presentation that can be adapted based on the brain state or stimulation response.

An important aspect of the method is its open-access availability. NeuroSimo, InVesalius neuronavigation, and robot control are accessible under open-source licenses. The method can be used with commercially available TMS devices connected to a robotic arm, with stimulation triggered by a LabJack USB peripheral. All features of the method except for stimulation location control can also be used for TMS without a robotic arm.

### 5.2. Technical considerations

EEG preprocessing and decoding with a pre-trained algorithm took, on average, 150 ms, and robotic arm movement from the vertex to the M1 target took, on average, less than 8 s. Although slight variations in latency from robotic arm movement to the pulse existed across targets, these variations demonstrated the robotic targeting system’s ability to reach the precise stimulation target even if the participant moved. The robotic arm velocity is also limited by safety thresholds. However, robotic TMS control makes stimulation completely automatic and thus removes the need for a human operator to move the coil. In protocols where mTMS is used and the stimulated targets are close enough to be stimulated without moving the mTMS coil set, there will be no delay due to robot movement. In such protocols, stimulus location and orientation can be changed electronically in less than a millisecond (Nieminen et al., 2019; Souza et al., 2022).

We recommend performing real-time EEG-trial filtering in a slightly different way than in the proof-of-concept measurement. High-pass filtering 2-s trials from 0.1 Hz may have caused edge effects in the slow-drift range in the whole trial. These drifts might have affected decoding performance, although they do not influence the frequencies of interest for ME (*ca.* 8–30 Hz).

The EEG-decoding algorithm exhibited excellent sensitivity but only moderate specificity for left-hand-movement trials. For the two other trial types, performance was lower. One reason that could affect the algorithm’s performance is the mTMS coil set moving between targets, since the pressure of the coil onto an EEG electrode might induce high-amplitude noise into the EEG channel, degrading the signal quality. This factor should be addressed when designing experiments with the method, especially when considering EEG preprocessing. The maximum coil-set pressure should be optimized to avoid such disturbances.

The decoding algorithm is not the focus of the present study, as the method is not limited to a single algorithm type. We demonstrated that a machine-learning algorithm can be trained for each participant by collecting the training data in the same session as the TMS experiment; NeuroSimo’s logging feature allows timestamping and storing trial types during the session for processing training data.

### 5.3. Applications

Our method enables controlling stimulation with volitional brain activity, making it ideal for protocols involving ME or MI. For example, the proof-of-concept experiment protocol can be directly adapted from using ME to MI. Thus, the method has many potential applications in motor rehabilitation with TMS, such as EEG-based BCI therapies in spinal cord injury (Shulga et al., 2026, 2021) and stroke patients (Liu et al., 2023; Pichiorri et al., 2015; Ramos-Murguialday et al., 2019). Combining the existing TMS rehabilitation protocol for spinal cord injuries with MI-BCI would automate the therapy and likely lead to faster rehabilitation timelines (Shulga et al., 2021; Royter and Gharabaghi, 2016). Stimulation during MI can inhibit functions in undesired locations (*e.g.*, inside tumors (Rivera-Rivera et al., 2017); prehabilitation) and guide functional reorganization by facilitating new cortical areas. The fully automatic operation in our method also paves the way for home-based or remote treatment.

Our method can additionally be helpful in developing brain-state-dependent depression treatment. Timing TMS to specific EEG events has been studied in patients with depression (Faller et al., 2022; George et al., 2023; Price et al., 2010; Zrenner et al., 2020). Our method enables new rehabilitation and treatment paradigms by adding the ability to adaptively optimize stimulation location and timing with complex brain states decoded from EEG, potentially improving treatment outcomes.

The method allows probing and studying causal interactions in network brain functions. For example, the motor network could be probed at different times of the movement planning–execution process in different network locations. As our method enables causal mapping, such measurements would offer unique insights into the dynamics of brain networks.

## 6. Conclusion

We demonstrate that both TMS location and timing can be controlled based on EEG-decoded brain state. The presented method enables a wide range of brain-state-dependent and closed-loop stimulation experiments and facilitates the development of neurological rehabilitation protocols. The software components of the method are openly available. The method allows control of brain stimulation using volitional brain activity, such as motor imagery. This paves the way for novel non-invasive personalized rehabilitation for conditions such as spinal cord injury.

## CRediT authorship contribution statement

**Renan H. Matsuda:** Conceptualization, software, methodology, investigation, data curation, formal analysis, visualization, project administration, funding acquisition, writing – original draft, writing – review and editing. **Matilda Makkonen:** Conceptualization, software, methodology, investigation, data curation, visualization, project administration, funding acquisition, writing – original draft, writing – review and editing. **Ivan Zubarev:** Methodology, software, writing – review and editing. **Olli-Pekka Kahilakoski:** Software, writing – review and editing. **Leevi A. Kinnunen:** Formal analysis, visualization, writing – review and editing. **Mila Nurminen:** Methodology, writing – review and editing. **Simo-Pekka Simonaho:** Resources, supervision, writing – review and editing. **Risto J. Ilmoniemi:** Supervision, funding acquisition, writing – review and editing. **Victor H. Souza:** Supervision, funding acquisition, writing – review and editing. **Pantelis Lioumis:** Conceptualization, methodology, supervision, project administration, funding acquisition, writing – review and editing.

## Data availability

The full data is available from the corresponding author on reasonable request.

## Declaration of generative AI and AI-assisted technologies in the writing process

During the preparation of this work the authors used ChatGPT-5.5 in order to edit and proofread the language of the raw text. After using this tool, the authors reviewed and edited the content as needed and take full responsibility for the content of the published article.

## Acknowledgments

This work was supported by Vaikuttavuussäätiö decision 366, Jenny and Antti Wihuri Foundation Grant 00250241 (RHM), Finnish Cultural Foundation (MM) under Grant 00250619, Emil Aaltonen Foundation (MM) under Grant 240110 K1, Research Council of Finland (Decision No. 349985), and the European Research Council (ERC Synergy) under the European Union’s Horizon 2020 research and innovation programme (ConnectToBrain; grant agreement No 810377).

